# ZipHiC: a novel Bayesian framework to identify enriched interactions and experimental biases in Hi-C data

**DOI:** 10.1101/2021.10.19.463680

**Authors:** Itunu G. Osuntoki, Andrew Harrison, Hongsheng Dai, Yanchun Bao, Nicolae Radu Zabet

## Abstract

Several computational and statistical methods have been developed to analyse data generated through the 3C-based methods, especially the Hi-C. Most of existing methods do not account for dependency in Hi-C data. Here, we present ZipHiC, a novel statistical method to explore Hi-C data focusing on detection of enriched contacts. ZipHiC implements a Bayesian method based on a hidden Markov random field (HMRF) model and the Approximate Bayesian Computation (ABC) to detect interactions in two-dimensional space based on Hi-C contact frequency matrix. ZipHiC uses data on the sources of biases related to contact frequency matrix, allows borrowing information from neighbours using the Potts model and improves computation speed by using the ABC model. In addition to outperforming existing tools on both simulated and real data, our model also provides insights into different sources of biases that affects Hi-C data. We show that some datasets display higher biases from DNA accessibility or Transposable Elements content. Furthermore, approximately half of the detected significant interactions connect promoters with other parts of the genome indicating a functional biological role. Finally, we found that the micro-C datasets display higher biases from DNA accessibility compared to a similar Hi-C experiment, but this can be corrected by ZipHiC.

## 1 Introduction

Distant regulatory elements and their target genes are often separated by large genomic distances. In order for the regulatory element to activate a target gene, they need to come in 3D proximity (Bonev and Cavalli, 2016; Hua *et al*., 2021). This indicates that the spatial organisation of the genome is intimately related to genome regulation and better understanding the 3D organisation of the genome is important to disentangle the contribution of different factors to gene regulation. One of the recently developed genome-wide proximity ligation assay is the Hi-C technique (Lieberman-Aiden *et al*., 2009), which is a chromosome conformation capture (3C)-based method. Hi-C is able to detect interactions (short-range and long-range) within and between chromosomes at high resolutions. While in mammalian system, resolutions of 5 *Kb* have been achieved (Rao *et al*., 2014), in smaller genomes, such as *Drosophila*, sub-kilobase pair resolutions were obtained from Hi-C experiments (Eagen *et al*., 2017; Cubenãs-Potts *et al*., 2017; Chathoth and Zabet, 2019). In addition, datasets generated by Hi-C are highly reproducible between replicates and often highly conserved between tissues (Ghavi-Helm *et al*., 2014). Recent technological advances have pushed the resolution of conformation capture methods to base pair resolution in mammalian systems (Hua *et al*., 2021).

The data generated by a Hi-C experiment can be represented as a matrix of contact frequencies between pairs of fragments along the genome. These matrices are associated with biases ((Yaffe and Tanay, 2011)), such as the restriction fragment length, GC-content of trimmed ligation junctions and mappability, but many additional factors may also contribute to the contact counts. Correcting for these biases is important and there has been several methods being proposed to take them into account (Yaffe and Tanay, 2011; Imakaev *et al*., 2012; Hu *et al*., 2013; Servant *et al*., 2015).

The Iterative correction and eigenvector decomposition(ICE) has been the most widely used method to account for biases associated with the Hi-C data, due to its simplicity and being parameter-free by assuming equal visibility across all fragments (Imakaev *et al*., 2012). This equal visibility assumption considers that all fragments can be probed by the method with same probability. However this assumption is not always true, because the visibility of areas could vary. In addition, ICE is computationally intensive because the Hi-C interaction matrix is of size *O*(N^2^), where N is the number of genomic regions.

The study of (Rao *et al*., 2014) generated one of the highest resolution map of the 3D organisation of the human genome by using a *in situ* Hi-C to probe the 3D architecture of genomes for DNA-DNA proximity ligation in intact nuclei. This has revealed that the human genome is organized into sub-compartments globally and contains about 10, 000 chromatin loops (Rao *et al*., 2014). To account for biases in Hi-C data, (Rao *et al*., 2014) adopts the matrix-balancing proposed in (Knight and Ruiz, 2013). In particular, peaks are called only when a pair of fragment shows elevated contact frequency relative to the local background; i.e., peaks are called when the peak pixel is enriched as compared to other pixels in its neighborhood.

Other methods take into account potential dependence among pairs of fragments (Jin *et al*., 2013). In order to accurately identify the chromatin interactions and loops with high sensitivity and resolution, they used data filtering technique based on the strand orientation of Hi-C pairedend reads. This also allows detection of short distance interactions between restriction fragments and their analysis shows the effects of GC-content and mappability on the observed contact frequency. Interestingly, there seems to be a linear relationship between average trans-contact frequency and mappability (Jin *et al*., 2013).

Neighbouring regions often interact with the same fragments suggesting that these anchors are part of a large region. Some of the existing methods are based on one-dimensional calling approaches, which do not consider useful information that can be gained using the two-dimensional approach. The first method to take into account the spatial dependency of Hi-C is the HMRFBayesHiC algorithm (Xu *et al*., 2016b). In particular, HMRFBayesHiC models the neighbouring fragments in the context of a two-dimensional contact matrix generated from Hi-C. This algorithm assumes that not all peaks will have similar strength and clustering patterns. Nevertheless, it also involves having prior information about the expected count frequency distribution to account for biases, which is often unknown. One of the biggest shortcomings of this approach is that it is computationally intensive and chromosome wide computations, even in smaller genomes, are not feasible.

FastHiC is a novel hidden Markov random field (HMRF)-based peak caller to detect long-range chromosomal interactions from Hi-C data (Xu *et al*., 2016a). The FastHiC method is based based on the HMRFBayesHiC (Xu *et al*., 2016b) and uses simulated field approximation, which approximates the joint distribution of the hidden peak status by a set of independent random variables. In particular, FastHiC approximates the Ising distribution by a set of independent random variables, enabling tractable computation of the normalising constant in the Ising model. Despite this improvement in computation time, FastHiC is still computational intensive and chromosome wide computations are still challenging.

Finally, all these previous methods, often classify the observations into only two classes: non-random contacts (peaks) and random contacts (noise). Nevertheless, it is possible to have more than two classes due to the nature of the Hi-C approach. For example, a non-random contact may have similar bias information to a random contact, which may lead to misclassification of this pair of fragments by existing method.

In this paper, we present ZipHiC, a hidden Markov random field based Bayesian approach to identify significant interactions in Hi-C data. This new model addresses several issues with current models. First, we improve on existing methods by introducing dependency of neighbouring fragments in the two-dimensional space and by adopting the Approximate Bayesian Approach (ABC) to deal with the intractable normalizing constant in the Potts model, a Markov random field-based model (Wu, 1982). Second, our model is computationally tractable and can be applied chromosome wide. Third, the number of classes under consideration can be naturally extended to more than two. We focus our analysis on intra-chromosomal interactions due to the fact that about 95% of non-random interactions are found within chromosomes (Jin *et al*., 2013; Xu *et al*., 2016b). Most importantly, we use ZipHiC to model Hi-C contact maps in *Drosophila* cells and human cells and explore biases introduced by GC content, transposable elements (TEs) and DNA accessibility. Finally, we also model micro-C data in human ES cells and compare it to a similar Hi-C dataset in terms of the identified significant contacts and biases.

## 2 Materials and Methods

### 2.1 ZipHiC

#### 2.1.1 Notations

ZipHiC uses the contact matrix between pairs of fragments generated from Hi-C experiments. Let *y*_*ij*_, 0 ≤ *i < j* ≤ *N* denote the observed contact frequency between fragment *i* and fragment *j* in *N* total fragments and *D*_*ij*_ represent the genomic distance between fragment *i* and fragment *j*. Let *GC*_*ij*_ represent the average percentage of Guanine and Cytosine, *TE*_*ij*_ represent the average number of transposable elements (TEs) and *ACC*_*ij*_ represent the average DNA accessibility score in fragments *i* and *j*. For simplicity, we use *s* = {*i, j*} to denote the interaction pair of fragments *i* and *j* and use *D*_*s*_, *GC*_*s*_, *ACC*_*s*_ and *TE*_*s*_ to denote the observation value for interaction *s*.

#### 2.1.2 Mixture model for data

We use the *K*-component mixture density to model our data *y*_*ij*_, where the first component is a zero-inflated Poisson (ZIP) distribution for noise (see below), while the other components follow Poisson distributions:

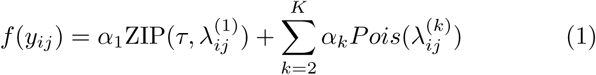

where *τ* is the probability of extra zeros, 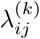 is the mean of the *k*th component. *α*_*k*_ is unknown percentage of *k*th component where 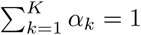.

The above mixture model can be interpreted via a latent variable framework. We introduce the latent variable *z*_*ij*_ = *k, k* = 1, 2, …, *K*, where *z*_*ij*_ = *k* means that *y*_*ij*_ follows the distribution of component *k*. Furthermore, 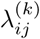 represents the mean interaction of fragments *i* and *j* if it is from the *k*th component. We propose increasing to *K*th components which makes the framework more flexible for different scenarios.

Due to the fact that the Hi-C contact map displays excess zero-counts and that the mean and variance are not the same, we assume that the noise follows a ZIP distribution rather than a Poisson distribution. In particular, a ZIP distribution has the mean (1−*τ*)*λ* and variance *λ*(1−*τ*)(1+*τλ*). Furthermore we assume that the sources of biases can be corrected by modeling 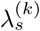 with *s* = {*i, j*}, *k* = 1, 2, · · ·, *K* as

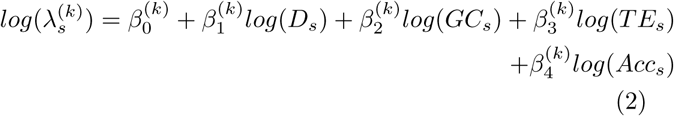

### 2.1.3 Potts Model

To introduce the spatial dependency, our method utilizes the HMRF for the hidden components. The HMRF is a generalization of the hidden Markov model (HMM). The HMRF has been widely used in areas such as image analysis ((Zhang *et al*., 2001)), gene expression data ((Wei *et al*., 2008)) and population genetics study ((François *et al*., 2006)). We adopt the Potts model ((Wu, 1982)) which is a Markov random field-based method that provides a flexible way to model spatially dependent data as our prior for the latent variable *z*_*s*_. The latent variable **z** adopting the Potts model is written as

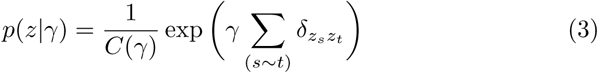

where 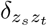 is the Kronecker symbol which takes the value 1 when *z*_*s*_ = *z*_*t*_ and 0 otherwise. Label *t* defines the neighboring fragment pairs of *s*, i.e. *s* ∼ *t* means *s* and *t* are neighbours in the Hi-C matrix. The set of latent variables *z*_*ij*_ are modelled as a 2-dimensional HMRF, so the latent variable *z*_*s*_ depends on the status of the neighbors of *s* = {*i, j*}, 𝒩_*s*_ = {(*i* + 1, *j*), (*i* − 1, *j*), (*i, j* + 1), (*i, j* − 1)}. The neighbouring 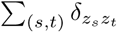 can be interpreted as the sum of the influence of neighbours of *s*. Here *γ* is a non-negative interaction parameter, with value 0 resulting in an independent uniform distribution on *z*_*ij*_. Larger values of *γ*, such as *γ* = 1, corresponds to a high level of spatial interaction, and the probability of pairs of neighbours being in the same component is very high. *C*(*γ*) is the normalizing constant, also known as the partition function, which is written as

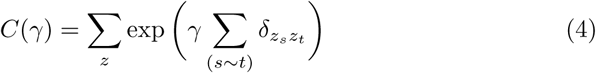

where ∑_*z*_ indicates the summation over *z*_*s*_ at all interactions *s* and it depends on the interaction parameter *γ*. The normalizing constant is computationally intractable in higher order. To overcome this complication, methods such as the likelihood-free approach can be used. Here we use the Approximate Bayesian Model (ABC) (Beaumont *et al*., 2002).

#### 2.1.4 Approximate Bayesian Model (ABC)

The ABC algorithm (Beaumont *et al*., 2002) used here can be described as follows:

- For a given dataset *Y* = (*y*_1_, *y*_2_, …, *y*_*n*_) that is associated with the models in equations (1), (2) and (3), simulate an initial value *γ*_0_ from the prior distribution *π*_0_(*γ*);
- Generate a parameter value from the posterior distribution *π*(*γ*|*Y*) ∝ *π*_0_(*γ*)*p*(*z*|*γ*);
- A new value of *γ*\* and *y*\* is simulated jointly from (1), (2) and (3);
- Compute the absolute distance or euclidean distance *d* between the simulated data and the observed data;
- fix a tolerance *ϵ* or use an empirical quantile of *d*(*y*\*, *y*) which often corresponds to 1% quantile ((Beaumont *et al*., 2002))
- Accept *γ*\* if the absolute distance is less than *ϵ*, otherwise reject and start from step 1 again.

#### 2.1.5 Bayesian Inference

In order to infer parameters, we adopt the Bayesian approach which is based on the posterior distribution. The posterior distribution is the product of the prior and likelihood. For our prior, we make use of the Empirical Bayes approach, which uses a hierarchical structure to determine the prior, where the prior is determined by a distribution with parameters called hyper-priors. The hyper-priors are estimated from the dataset which means that it is less affected by mis-specification of priors.

We also use the conventional Bayesian approach. For the conventional Bayesian approach, we set the priors of our *β*s to follow the normal distribution. For example, we set the prior of 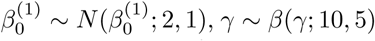 and set *π*_0_ = 0.6. To estimate the impact of priors, we consider other priors; see *Results* and *Supplementary Material*.

### 2.2 Datasets and preprocessing

#### 2.2.1 Drosophila dataset

To test the performance of the model, we used a high resolution Hi-C map of Kc167 cell lines in *Drosophila* from (Eagen *et al*., 2017). The raw data was downloaded and preprocessed with HiCExplorer following the set of parameters from (Chathoth and Zabet, 2019). Briefly, we aligned each pair of the PE reads to *Drosophila melanogaster* (dm6) genome (dos Santos *et al*., 2015) using BWA-mem (Li and Durbin, 2010) (with options -t 20 -A1 -B4 -E50 - L0). HiCExplorer was used to build and correct the contact matrices and detect enriched contacts (Ramirez *et al*., 2018). The contact matrices were built using 2 Kb bins and then exported in text format to be loaded in R.

For DNA accessibility in *Drosophila* Kc167 cells data we used DNaseI-seq data from (Kharchenko *et al*., 2010), while, for TE annotation in *Drosophila*, we used FlyBase (dos Santos *et al*., 2015).

We detected TADs using HiCExplorer at 2Kb resolution, similarly as done in (Chathoth and Zabet, 2019). Briefly, TADs had at least 20 Kb width, a P-value threshold of 0.01, a minimum threshold of the difference between the TAD-separation score of 0.04, and FDR correction for multiple testing (–step 2000, –minBoundaryDistance 20000 –pvalue 0.01 –delta 0.04 – correctForMultipleTesting fdr).

#### 2.2.2 Human datasets

We also used Hi-C and micro-C datasets in H1-hES cells from (Krietenstein *et al*., 2020). We used the same preprocessing pipeline as for the *Drosophila* dataset. Briefly, we aligned each pair to the human genome hg38 (Schneider *et al*., 2017) using BWA-mem (Li and Durbin, 2010). HiCExplorer was used to build and correct the contact matrices at 10 Kb resolution and detect enriched contacts (Ramirez *et al*., 2018).

Furthermore, we used DNaseI-seq for DNA accessibility from ENCODE consortium (Thurman *et al*., 2012) and TE annotation from RepeatMasker (Smit, 2015).

### 2.3 Comparison to other tools

In this manuscript, we compare our new method ZipHiC to three other tools: *(i)* FastHiC (Xu *et al*., 2016a), *(ii)* HiCExplorer (Ramirez *et al*., 2018) and *(ii)* Juicer (Durand *et al*., 2017). First, we generated the enriched interactions using a JAVA implementation of FastHiC which use excepted counts and, for that, we the values estimated by the HiCExplorer (Ramirez *et al*., 2018).

Second, we used the HiCExplorer generated matrices and corrected them using the following values: (*i*) [−1.8, 5.0] for Hi-C in Kc167 cells, (*ii*) [−2.4, 5.0] for Hi-C in H1-hES cells and (*iii*) [−2.0, 5.0] for micro-C in H1-hES cells; see Supplementary Figure S1 (Ramirez *et al*., 2018).Then, we generated the enriched contacts from the corrected matrix using `hicFindEnrichedContacts` tool with observed over expected method (`--method obs/exp`) (Ramirez *et al*., 2018).

Third, we used Juicer to generate enriched contacts by calling `dump` tool from Juicer tools. In particular, we used the observed over expected method (`oe`) and Knight-Ruiz normalisation (`KR`) at 2 *Kb* resolution for the Hi-C data in Kc167 cells and at at 10 *Kb* resolution for the Hi-C and micro-C data in H1-hES cells (Durand *et al*., 2017).

The R scripts used to perform the analysis can be downloaded from https://github.com/igosungithub/HMRFHiC.git.

## 3 Results

### 3.1 Using the two component model on simulated data

First, we consider the case of a two component model (signal and noise) and evaluated whether this model can correctly estimate the sources of biases associated with Hi-C contact matrix using simulated data. We simulated a dataset of *n* = 2, 500 observations from the mixture model (1), with *K* = 2. The simulation studies are based on outputs of MCMC algorithms with 20, 000 iterations and 10, 000 burn-in steps. We considered using either informative prior or Empirical Bayes method, which has been used previously to analyse missing data (Carlin and Louis, 2000). Furthermore, there are three cases under different component proportions: *(i)* when the proportion of the noise is greater than the signal, *(ii)* when the proportion of the noise and the signal is the same, *(iii)* when the proportion of noise is less than the signal. Finally, we also used different starting values to justify the convergence of MCMC algorithms.

We study the sensitivity of our model to different set of prior parameters values using the traditional informative prior and Empirical Bayes method. The latter, the prior of Empirical Bayes method, is based on the hyper-prior determined by the dataset. Table S1 shows that the two component model are able to estimate the true value accurately when we using either the Informative or the Empirical Bayes method for the prior distribution. In order to illustrate the effect of using one of the priors (fixed prior or Empirical Bayes), we included only one covariate, *D*_*ij*_ (genomic distance) from 2. Our results show that the estimates of the posterior means of the parameters are accurate for both approaches to infer prior distribution. For our downstream analysis, we use the Empirical Bayes method.

Next, we evaluated the estimated posterior means of the parameters for our regression model (see equation 2). We used fixed informative prior and the component percentages (*α*s) in equation 1 are set as *α*_1_ = 0.7 and *α*_2_ = 0.3, showing a higher percentage of noise to signal. Table S2 shows that our method was able to estimate the true parameters accurately despite the higher noise. We also check our estimated posterior means with respect to their credible intervals, which are usually used in Bayesian analysis and have similar interpretation to confidence intervals. The main differences between our estimated posterior means fall within ±0.02 of the true values and our estimated posterior means are all significant as they fall within the 90% credible intervals. In addition, when evaluating Tables S1 and S2 and analysing the trace plots of all our simulations, we did not observe label switching; i.e., we are able to identify each components parameters distinctly without any unidentifiability issues. Furthermore, in Tables S3 and S4, we show that our method is also robust to different proportions of noise and signal (see *Supplementary Material*).

### 3.2 Hi-C Data analysis with a two components model

Following the validation of our model on simulated data, we next use the two component ZipHiC model on real Hi-C data. In particular, we use a dataset from (Eagen *et al*., 2017) in Kc167 cell line in *Drosophila* at 2 *Kb* resolution and focus this analysis on chromosome 2L. As mentioned earlier, the aim of our proposed method is to detect significant interactions, which we called true signal, by taking into consideration the biases associated with Hi-C dataset.

First, we considered the 31, 375 observations from a 82 *Kb* fragment (region 2L:1-82000), resulting in 250 unique pair of fragments in order to compare our method to existing statistical methods. FastHic (Xu *et al*., 2016a) is an updated version of the HMRFBayesHiC (Xu *et al*., 2016b) as both methods use a hidden Markov random field (HMRF) based Bayesian method and Ising model (Ising, 1925), which accounts for spatial dependence for peak calling. Note that, we only used 31, 375 observations, because of the high computation time of the FastHic (Xu *et al*., 2016a). In contrast to ZipHiC, FastHic (Xu *et al*., 2016a) method involves calculating the time expected frequencies, which is computation intensive and can be done using the approach in (Lieberman-Aiden *et al*., 2009).

Based on the Monte Carlo draws from the posterior distribution of our ZipHiC model, we can compute whether the estimated values of our parameters are significant or not (see posterior means values in Tables S5 and S6 in *Supplementary Material*). Figure 1 shows the Venn diagram of the biologically significant interacting pairs of fragments using ZipHiC two component model compared to FastHic (Xu *et al*., 2016a). ZipHiC recovers 87% (21, 061) of the interactions detected by FastHic (Xu *et al*., 2016a); see Figure 1. We notice that the FastHic (Xu *et al*., 2016a) method discovered additional 3, 106 pairing fragments as being biologically significant and this suggests that our model is slightly more conservative in detecting significant interactions. Interestingly, both methods detected 7, 134 pairing fragments as noise (random collision). Further investigation of the additional significant pairing fragments detected by the FastHic (Xu *et al*., 2016a) and not by our method, shows that the FastHic (Xu *et al*., 2016a) has higher false discovery rate than our method by falsely classifying the pairing fragments with 0 frequency as being significant.

**Figure 1:**
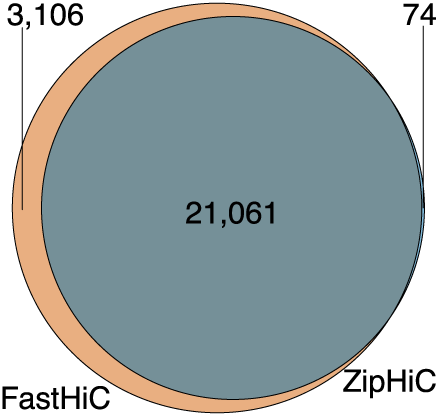
Comparison between ZipHiC and FastHiC. Venn Diagram showing true signal comparison between our proposed method (ZipHiC) and FastHiC on sub region of chromosome 2L in *Drosophila* Kc167 cells.

### 3.3 Hi-C Data analysis with a three components model

One limitation of previous studies was the limitation to two components (noise and signal). Here, we further increase the number of components from *K* = 2 to *K* = 3 by adding a new component. This new component accounts for interacting pairs of fragments that ZipHiC has misclassified as signal due to conflicting information both in the contact frequencies and sources of bias and, thus, we call this new component *false signal*. For example, if a pair of interacting fragment have high contact frequency but their sources of bias closely exhibits that of the noise component, this pair of fragment can be classified to the false signal component.

First, we compared the detected significant interactions in the three component ZipHiC model with the ones in the two component one and from FastHic. Figure 2 shows that by adding an additional component, we detect less than 1% of additional pairs of fragments (231) overlapping with the FastHic ((Xu *et al*., 2016a)) method.

**Figure 2:**
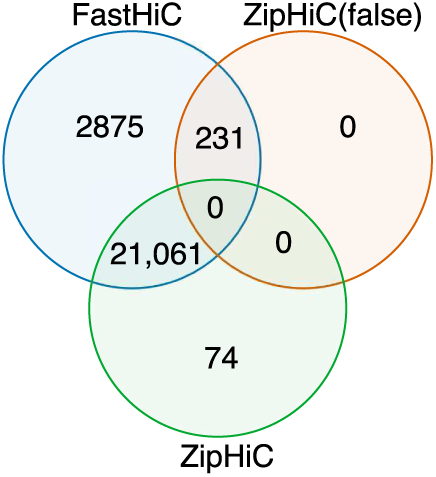
Venn Diagram showing comparison between the HMRF (Xu *et al*., 2016b), ZipHiC-2 (our true signal) and ZipHiC-3 (our false signal) of the sub region of Chromosome 2L of *Drosophila Melanogaster*.

To evaluate whether the new component in our method (false signal) results in better performance of our method, we conducted model selection analysis using the Deviance Information Criterion (Spiegelhalter *et al*., 2002) and in particular, we used a modified DIC method (Li *et al*., 2020) for latent variable models. The value of the DIC for two component is −331, 344, 746 and for the three component is −401, 662, 547. These results show that the best model to analyse this particular Hi-C dataset is the three components model (thus, including the false signal).

To better understand the contributions of the different components, we investigated the posterior means of our estimated *β*s for the noise, signal and false signal components (see Table 1). The values of *β*s correspond to the coefficients of the intercept and the log of distance, GC-content, TEs content and DNA accessibility. The posterior means of noise levels of the interaction for all components, except GC content, have *β* values with negative signs, indicating the noise and signal are negatively correlated. The negative sign of *β*_1_ parameter (distance) indicates that when distance between two fragments increases the average of their interaction noise decreases. Similarly, for *β*_3_ (TEs) and *β*_4_ (DNA accessibility), our results indicate that the higher the TEs content or the level of DNA accessibility is the lower the interaction noise will be, but only for DNA accessibility the effect is large. In other words, noise levels in the Hi-C signals are higher in dense chromatin and will have a higher impact on the observed enriched interactions, unless correctly accounted for. Nevertheless, for *β*_2_ (GC content), we found that the GC-content increase corresponds to an increase in the interaction noise, but, while this is significant, the contribution of GC content is relatively small to the noise levels in Hi-C data. Interestingly, we noticed from Table S6 and Table 1 that our estimated posterior means for the noise components are similar if we use a two component or a three component model. This can be seen as most of the third component (false signal) in our model is influenced by the second component (true signal).

**Table 1:** Posterior means of our estimated *β*s as shown in equation 2 for noise, signal and false signal components. The 95% credible intervals are shown inside the brackets. For the 90% credible intervals, see Table S7 in *Supplementary Material*.

| Parameters | Posterior mean (noise) | Posterior mean (signal) | Posterior mean (false signal) |
| --- | --- | --- | --- |
| $\beta_0$ (intercept) | -84.00 (-84.90, -83.64) | 13.06 (12.80, 13.35) | 499.34 (498.39, 500.21) |
| $\beta_1$ (distance) | -10.05 (-10.16, -10.03) | -0.90 (-0.92, -0.89) | -64.16 (-64.45, -63.97) |
| $\beta_2$ (GC content) | 0.34 (0.34, 0.35) | 0.36 (0.35, 0.37) | 0.30 (0.09, 0.57) |
| $\beta_3$ (TEs) | -0.76 (-0.79, -0.68) | -0.10 (-0.16, -0.03) | 0.54 (-0.40, 1.03) |
| $\beta_4$ (Accessibility) | -3.54 (-3.57, -3.44) | 0.15 (0.11, 0.20) | -0.70 (-1.04, -0.15) |

Next, we observed that the posterior mean of our false signal intercept is higher than that of the noise and signal components, which implies what the value of the log of our *λ* in (2) will be if all the sources of bias are kept at 0. For the false signal component, we noticed that the posterior mean and credible intervals for distance (*β*_1_) parameter of the false signal component is significant and the negative value indicates that as the increase distance of two fragments results in a decrease in the false signal interaction. The effect size of distance on false signal is higher than compared to noise and was previously unaccounted for. For DNA accessibility (*β*_4_), the negative value of the posterior mean and the credible intervals means that increase DNA accessibility leads to a decrease in the false signal interaction, but this is relatively small. Similarly for the posterior mean of the GC content (*β*_2_), the value is positive and indicates that as higher GC content corresponds to an increase in the false signal. However for TEs (*β*_3_) the credible intervals of false signal component covers 0, which means the result is not significant.

Furthermore, we noticed that the posterior mean of true signal for GC content (*β*_2_) decreased when the third component (false signal) was added (compare from Tables 1 and S6). This means that the influence of GC content was reduced when taking into account false signal. In addition, we noticed that the estimated posterior mean of (TEs) *β*_3_ for the signal component is significant and the false signal component is insignificant when the third component was added. This indicates that in order to properly estimate the true signal over TEs a three component model might be required and previous models that did not include a false signal might have obtained inaccurate enriched contacts over TEs.

### 3.4 Whole chromosome analysis using the three component ZipHiC model

Given that our model performs best with three components on this particular Hi-C dataset in *Drosophila* Kc167 cells, we analyse the whole of chromosome 2L using the three component ZipHiC model and identified 12.82M significant interactions. We observe that most of the detected significant interactions are found closer to the diagonal and that the significant interactions formed triangular shapes along the diagonal which sometimes overlap each others; see Figure 3.

**Figure 3:**
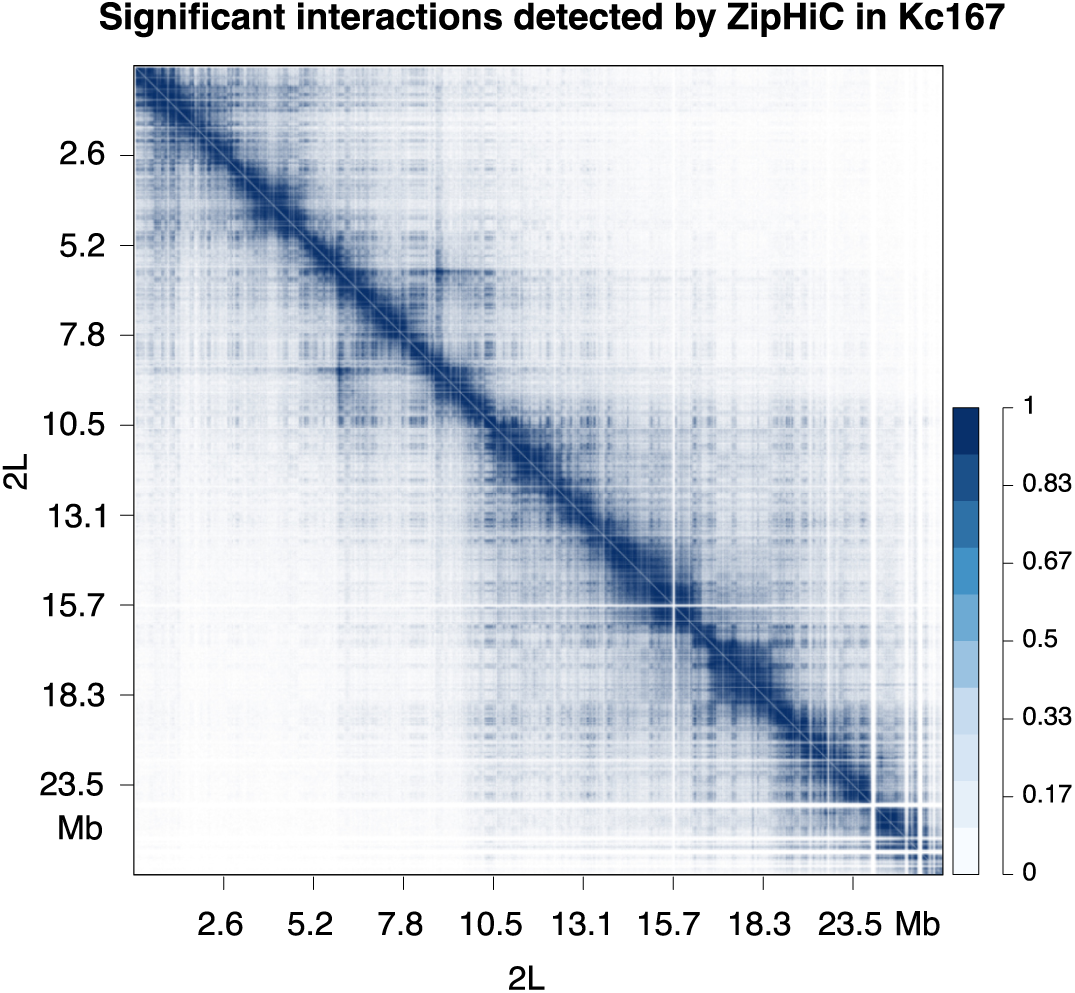
Significant interactions on chromosome 2L in Drosophila Kc167 cells. Heatmap showing significant interactions on chromosome 2L of *Drosophila* Kc167 cell line using ZipHiC three component model. The intensity of the colour indicates the probability, with darker colours representing higher probability.

These triangular shapes are Topologically Associated Domains (TADs) (Nora *et al*., 2012; Dixon *et al*., 2012; Sexton *et al*., 2012; Hansen *et al*., 2018) and are one of the main features of Hi-C data. However, we found that majority of significant interactions connect fragments that are very far apart (between 1 Mb and 10 Mb) (see Figure 4A), which are distances larger than the usual size of TADs in *Drosophila* (Ramirez *et al*., 2018; Chathoth and Zabet, 2019) and suggests that they connect fragments located in different TADs. Indeed, this is the case and approximately 98% of significant interactions are outside TADs (see Figure 4B). Interestingly, we found that almost half of the significant interactions connect promoters with other parts of the genome or with other promoters, which indicates they have a functional role (see Figure 4C). The majority of the significant interactions connect genes with either themselves or other genes, promoters or other regions of the genome (potentially enhancers). Note that we also performed a genome wide analysis and these results are true for all chromosomes (see Figure S2).

**Figure 4:**
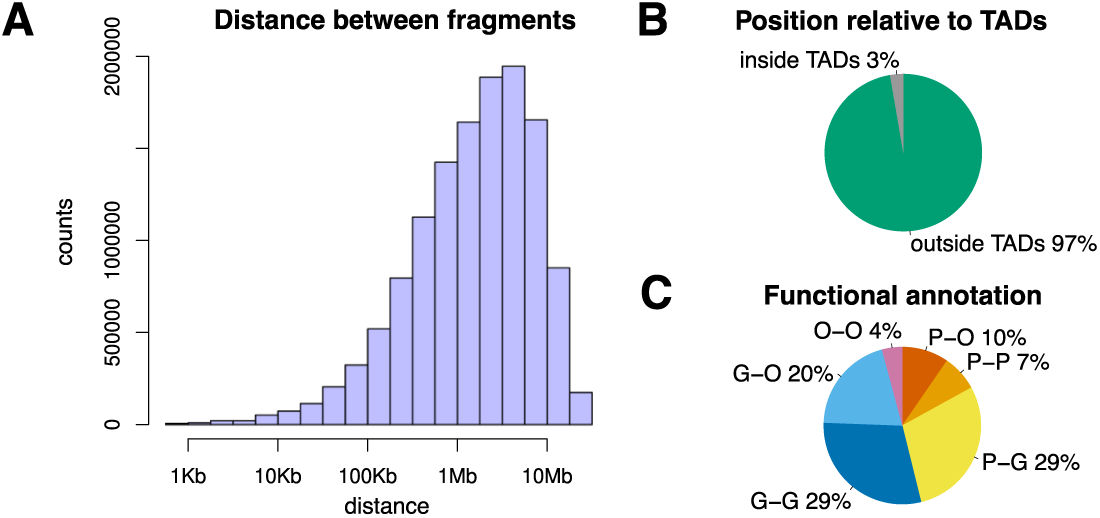
Characterisation of significant interactions on chromosome 2L in Drosophila Kc167 cells. (A) Distribution of the distance between the two fragments for all significant interactions. (B) classification of significant interactions as either outside TADs when the two fragments are located in different TADs or inside TADs when the two fragments are located in the same TAD. (C) Percentage of significant interactions that have promoters at one of the fragments. We consider the cases of: (P) promoters (200 bp upstream and 50 bp downstream of TSS), (G) genes (including exons, introns, 5’UTRs and 3’ UTRs and excluding promoters) and (O) other regions (excluding promoters and genes).

Finally, we compared the significant interaction detected by ZipHiC with significant interactions detected by two popular tools: HiCExplorer (Ramirez *et al*., 2018) and Juicer (Durand *et al*., 2017). Figure 5A shows that approximately half of the ZipHiC interactions are common with both HiCExplorer and Juicer (6.31M). In addition, ZipHiC detects 6.37M interactions detected only by HiCExplorer and missed by Juicer and 41K significant interactions detected only by Juicer and missed by HiCExplorer. ZipHiC distinctly identifies 129K significant interactions, which are missed by the other tools. Overall, we found that ZipHiC recovers almost all HiCExplorer (12.68M) significant interactions (98.9% overlap), but also an additional 170K significant interactions missed by HiCExplorer. Juicer seems to have a smaller overlap and only retrieves approximately half of the ZipHiC and HiCExplorer significant interactions.

**Figure 5:**
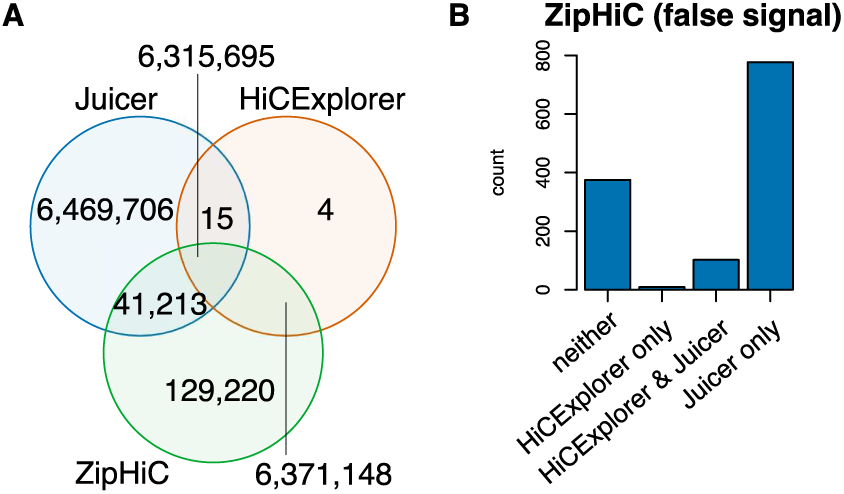
Comparison with other tools. (A) Venn Diagram showing the comparison between significant interactions detected by ZipHiC, HiCExplorer and Juicer. We analysed chromosome 2L in *Drosophila* Kc167 cells. (B) The number of false signals identified by ZipHiC detected as true signals by HiCExplorer and Juicer

Figure 5B shows the Venn diagram of the three component ZipHiC model with other methods by adding the false signal component. The highest number of false signals (245) overlap with the significant interactions detected by Juicer, further supporting the fact that this tool is less accurate in detecting significant interactions

8 pairing fragments classified by our method to be false signal overlaps with both HiCExplorer (Wolff *et al*., 2018) and Juicer (Durand *et al*., 2017). False signals identified by ZipHiC has 2 pairing fragments overlapping only with the HiCExplorer (Wolff *et al*., 2018). The false signal that does not overlap with any of the tools (116), indicates that while HiCExplorer can remove false significant interactions (only 10 false signals overlap with HiCExplorer), Juicer is more affected by these false significant interactions (253 false signals overlap with Juicer).

### 3.5 Analysis of micro-C data in human ES cells

Micro-C is a new and improved variation of Hi-C that can generate sub-kilobase pair 3D contacts map in mammalian systems (Hsieh *et al*., 2015; Krietenstein *et al*., 2020). To evaluate the capacity of ZipHiC to analyse micro-C data, we consider a small region on human chromosome 8 (60-70Mb) for which both micro-C and Hi-C data is available in human ES cells (Krietenstein *et al*., 2020). As we did previously, we consider both a two components and a three components model (*K* = 2 and *K* = 3) and use the DIC to select the best performing model (for the 3 components models of Hi-C and micro-C data see Table S8 and Table S9 respectively). Interestingly, in the case of this specific region on the human chromosome 8, the two component model has the lowest DIC (*DIC*_2_ = 194, 721.1 and *DIC*_3_ = 469, 950.5) and, thus, was selected for the analysis. This indicates that the human ES cell Hi-C and micro-C data in this region of the genome is not affected by false positive signals as it was the case with the *Drosophila* whole genome analysis in Kc167 cells.

Figure 6 shows the overlap of significant interactions identified by ZipHiC on micro-C and Hi-C datasets for this particular region of the human genome (60-70Mb). The Venn diagram shows that 96% (18, 498) of the significant interactions in the Hi-C dataset are recovered as significant interactions in the micro-C dataset as well and only a negligible number of interactions are missed (4%). Similarly, only 3% of the micro-C interactions are novel and previously missed by Hi-C. Our results confirm that micro-C can reproduce accurately the results of Hi-C despite a significantly lower library size.

**Figure 6:**
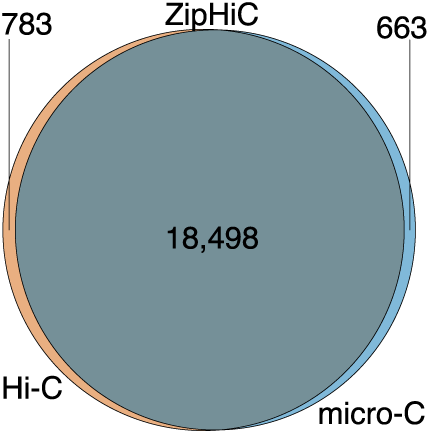
Venn Diagram showing significant interactions (signal) comparison identified by ZipHiC on micro-C and Hi-C data in human ES cells within 60-70Mb region of human chromosome 8.

We also investigated the overlap between the significant interactions identified by ZipHiC, Juicer and HiCExplorer and found that the three methods agree well (see Figure S3). Nevertheless, ZipHiC was also able to analyse the models and extract the sources of bias in the Hi-C and micro-C datasets. In micro-C, the chromatin is fragmented to mononucleosomes using micro-Coccal nuclease (MNase), which increases fragment density. The digestion with MNase raises the possibility that micro-C data is affected by DNA accessibility biases, which would not the case with Hi-C data.

Table 2 shows the model parameters for the two component model for both micro-C and Hi-C data. Interestingly, we observed an effect of the DNA accessibility on mean signal that is higher even compared to the effect of the distance between the fragments. Similar effect in the mean signal was also observed in the case of Hi-C data, but that was approximately half compared to the level observed in the micro-C data. In the case of the whole genome Hi-C analysis in *Drosophila*, we identified limited effects of accessibility on the mean signal but strong effects on the noise component. For this particular region in the human genome, we observed the opposite, strong biases introduced by accessibility in the mean signal (especially in the micro-C data), but significantly reduced biases on the noise of the signal. The beta values have positive sign indicating that more accessible regions of the genome display higher signal, but only modest biases in the noise levels.

**Table 2:** Posterior means of our estimated *β*s as shown in equation 2 for noise and signal components of human Chromosome 8, region 60, 000, 000 : 70, 000, 000 for data generated using the Hi-C and micro-C method. The 95% credible intervals are shown inside the brackets.

| Parameters (Hi-C) | Posterior mean (noise) | Posterior mean (signal) |
| --- | --- | --- |
| $\beta_0$ (intercept) | 0.88 (0.53, 1.36) | 11.13 (10.92, 11.36) |
| $\beta_1$ (distance) | 0.13 (-0.02, 0.25) | -0.79 (-0.81, -0.77) |
| $\beta_2$ (GC-content) | 0.33 (0.32, 0.33) | 0.32 (0.32, 0.34) |
| $\beta_3$ (TEs) | 1.01 (0.99, 1.15) | 0.02 (0.01, 0.03) |
| $\beta_4$ (Accessibility) | 0.50 (0.43, 0.59) | 1.00 (0.99, 1.03) |
| Parameters (micro-C) | Posterior mean (noise) | Posterior mean (signal) |
| $\beta_0$ (intercept) | 1.05 (0.81, 1.30) | 8.08 (7.65, 8.39) |
| $\beta_1$ (distance) | 0.14 (0.12, 0.17) | -1.41 (-1.42, -1.38) |
| $\beta_2$ (GC-content) | 0.33 (0.32, 0.34) | 1.02 (0.12, 1.80) |
| $\beta_3$ (TEs) | 10.00 (9.99, 10.02) | -0.37 (-0.41, -0.33) |
| $\beta_4$ (Accessibility) | 0.40 (0.35, 0.41) | 1.83 (1.70, 1.92) |

Furthermore, we also identified a strong contribution to the noise of the signal from the TE composition, specifically in the micro-C dataset, but also present in the Hi-C data despite being ten times lower. This means that a higher TE content leads to a higher noise, specifically in the micro-C data. In addition, micro-C data also display low bias of TE content in the mean signal, indicating that higher TE content leads to a slightly lower signal in micro-C, but not in Hi-C. Note that that in the case of whole genome analysis in *Drosophila*, there was only a relatively medium bias from TE content in the noise and false signal components, but not in the true signal one.

## 4 Discussion

In this manuscript, we introduce a new method called ZipHiC to analyse Hi-C and micro-C data. ZipHiC models the contact frequencies as a Zero-Inflated Poisson distribution due to the fact that this allows to model the presence of overdispersion which affects Hi-C data (Varoquaux *et al*., 2021). In addition, ZipHiC also uses a hidden Markov Random Field (HMRF) based Bayesian method, the Potts model, to help account for dependency in Hi-C dataset. Most importantly, the Potts model alows to increase the number of components (*k* = 2, 3, … *K*) and, thus, to account for additional components such as false signal. Finally, our method uses a likelihood free approach, ABC, to account for the limitation in the normalizing constant in the Potts model. Through our extensive simulations on simulated and real data, we show that our method outperforms existing methods in distinguishing between noise and signal.

First, we found that a three component model (specifically considering the false signal) performed better on a very high resolution dataset in *Drosophila* Kc167 cells (Eagen *et al*., 2017). However, a two component model (considering only the noise and the signal) performed best for the Hi-C and micro-C datasets in human ES cells (Krietenstein *et al*., 2020) on a region on chromosome 8. This indicates that the choice of whether to use a two component or a three component model needs to be driven by the data, since not all datasets will be affected by a false signal component.

In *Drosophila*, we found that the distance between fragments has the highest contribution to both the noise and the false signal, where interactions further from the diagonal displayes less noise and fewer false signals compared to interactions closer to the diagonal. DNA accessibility contributed strongly to the noise component and partially to the false signal in *Drosophila*. In particular, less accessible regions of the genome displayed higher noise and more false signals. We also observed a moderate effect of TEs on the noise component and false signal in *Drosophila*, where regions with higher content of TEs displayed lower noise, but higher false signals.

Majority of these significant interactions connect fragments that are located in different TADs and this is explained by the larger distance between the two fragments detected by ZipHiC in this dataset. The distance between fragments is larger than previously reported in *Drosophila* cells (Chathoth and Zabet, 2019), due to the fact that in this study we used a 2 Kb resolution and in the previous study a higher resolution was used (DpnII restriction sites, on average every 529 bp).

Most importantly, we identified that approximately half of these significant interactions in *Drosophila* connect promoters with either other promoters, genes or other regions of the genome. This raises the possibility that these significant interactions connect promoters with regulatory regions. Nevertheless, the large number of detected significant interactions and the number of enhancers identified in *Drosophila* cells (Arnold *et al*., 2013; Yanez-Cuna *et al*., 2014), indicate that most of them would not connect promoters with enhancers. This is likely the case and one possibility is that a large part of the significant interactions account for gene domains being formed at actively transcribed genes, where the promoter of the gene makes 3D contacts with different parts of the gene (exons, introns or 3’UTRs) (Rowley *et al*., 2019). Indeed, we found that majority of significant interactions involve genes, further supporting this model.

Furthermore, we found that micro-C data reproduces majority of the same significant interactions (96%) as a much larger Hi-C library. However, the micro-C data displays a higher bias in the signal to DNA accessibility (more accessible regions of the genome will display higher signals) even compared to distance between the fragments and this needs to be accounted for. Interestingly, in this particular region, the noise component was particularly affected by the TE content where more TEs lead to a higher noise in the micro-C data. The stronger effect of TEs on micro-C data in human cells is not surprising given the fact that human genome has a higher percentage of TEs compared to *Drosophila*.

A limitation to ZipHiC resides in the computation time when analysing the whole genomes. In the case of a standard computer with 4 cores, it takes more than 72 hours to analyse a whole genome dataset in *Drosophila* at 2 Kb resolution.

## Acknowledgements

We thank Liudmila A Mikheeva for useful comments and discussion on this work, Olivia Grant for comments on the manuscript and Stuart Newman for his support with the cluster.

