## Supplementary Material for "ZipHiC: a novel Bayesian framework to identify enriched interactions and experimental biases in Hi-C data"

### S1 Supplementary Figures

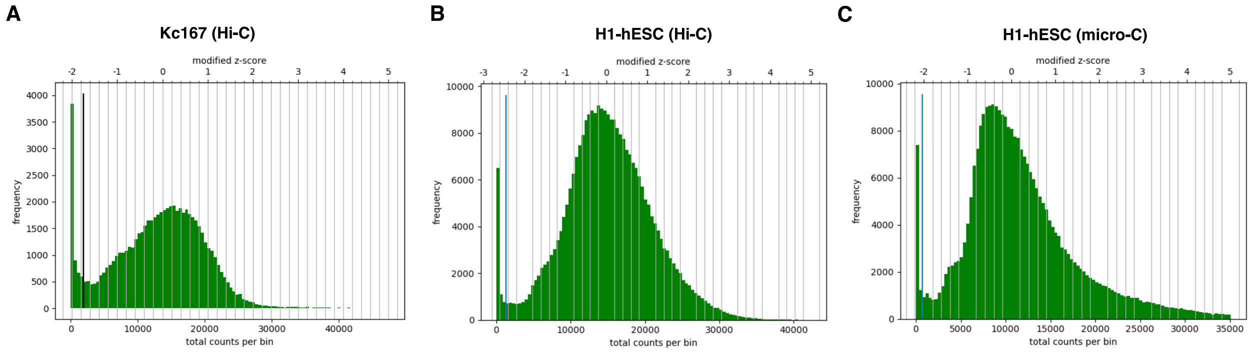


Figure S1: *Diagnostic plots for correction of Hi-C plots from HiCExplorer.* Histograms of the sum of contacts per bin. The vertical black line represents the lower threshold for removing bins with lower number of reads. We plotted the histograms for the three datasets used in this study: (A) Hi-C in Kc167, (B) Hi-C in H1-hES cells and (C) micro-C in H1-hES cells.

### S2 Model for Data

The complete likelihood function of the unknown parameters $(\boldsymbol{\beta}, \boldsymbol{\alpha}, \boldsymbol{\tau})$ given the data $y$ given $z$ can then be written as

$$l(\vec{\boldsymbol{\beta}}, \vec{\boldsymbol{\alpha}}, \boldsymbol{\tau}|\boldsymbol{y}, \boldsymbol{z}) \propto \prod_{i=1}^{n}\prod_{j=1}^{n}\left\{\left[\alpha_1\left(\tau + (1-\tau)e^{-\lambda_{ij}^{(1)}} + (1-\tau)\frac{\left(\lambda_{ij}^{(1)}\right)^{y_{ij}} e^{-\lambda_{ij}^{(1)}}}{y_{ij}!}\right)\right]^{I(z_{ij}=1)} \prod_{k=2}^{K}\left[(\alpha_k)\frac{\left(\lambda_{ij}^{(k)}\right)^{y_{ij}} \exp\left(-\lambda_{ij}^{(k)}\right)}{y_{ij}!}\right]^{I(z_{ij}=k)}\right\} \quad \text{(S1)}$$

The full posterior of $z$, $\vec{\boldsymbol{\beta}}$ and $\gamma$ given $y_{ij}$ is

$$Pr(\boldsymbol{z}, \vec{\boldsymbol{\beta}}, \gamma|\boldsymbol{y}) \propto l(\vec{\boldsymbol{\beta}}, \vec{\boldsymbol{\alpha}}, \boldsymbol{\tau}|\boldsymbol{y}, \boldsymbol{z}) l(\boldsymbol{z}|\gamma)\pi_0(\gamma)\tau_0(\vec{\boldsymbol{\beta}}) \quad \text{(S2)}$$

where $l(\boldsymbol{z}|\gamma) = \frac{e^{\gamma \sum_{s\sim t} \delta_{z_s z_t}}}{\sum_{z_s} e^{\gamma \sum_{s\sim t} \delta_{z_s z_t}}}$ is the Potts model, $s$ is fragment pair $i$ and $j$, and $t$ is the neighbours set of $s$; $(i-1, i+1, j-1, j+1)$.

In order to analyse our data and estimate our parameters, we make use of the Metropolis-within-Gibbs sampler and the Approximate Bayesian Computation (ABC), so the conditional posterior densities are needed.

#### S2.1 Conditional Posterior Density

The conditional posterior of $\tau$ is given as

$$Pr(\tau|\vec{\boldsymbol{\beta}}^{\mathbf{1}}, \boldsymbol{y}, \boldsymbol{z}) \propto \prod_{i=1}^{n}\prod_{j=1}^{n}[f^{(1)}(y_{ij}; \boldsymbol{\beta}^{(1)})]^{I[z_{ij}=1]} \cdot \pi_0(\tau) \quad \text{(S3)}$$

The conditional posterior of $\boldsymbol{\beta}$ is given by

$$Pr(\vec{\boldsymbol{\beta}}^{(\boldsymbol{k})}|\boldsymbol{y}, \boldsymbol{z}) \propto \prod_{i=1}^{n}\prod_{j=1}^{n}\prod_{k}^{K}[f^{(k)}(y_{ij}; \boldsymbol{\beta}^{(k)})]^{I[z_{ij}=k]}\pi_0(\boldsymbol{\beta}^{(k)}) \quad \text{(S4)}$$

Based on the definitions of $f^{(k)}(y_{ij})$ and equation S4, the conditional posterior of $\boldsymbol{\beta}^{(1)}$ for the noise component can be rewritten as

$$Pr(\vec{\boldsymbol{\beta}}^{(\mathbf{1})}|\boldsymbol{y}, \boldsymbol{z}) \propto \prod_{i=1}^{n}\prod_{j=1}^{n}\left[\left(\tau + (1-\tau)e^{-\lambda_{ij}^{(1)}} + (1-\tau)\frac{\left(\lambda_{ij}^{(1)}\right)^{y_{ij}} e^{-\lambda_{ij}^{(1)}}}{y_{ij}!}\right)\right]^{I(z_{ij}=1)} \exp\left\{-\frac{(\beta^{(1)} - m^{(1)})^2}{2(\sigma^2)^{(1)}}\right\} \quad \text{(S5)}$$

For the signal component, the conditional posterior of $\boldsymbol{\beta^{(k)}}$ based on definition of $f^{(k)}(y_{ij})$ and equation S4 can be rewritten as

$$Pr(\vec{\boldsymbol{\beta}}^{(\boldsymbol{k})}|\boldsymbol{y},\boldsymbol{z}) \propto \prod_{i=1}^{n}\prod_{j=1}^{n}\Big[\frac{\left(\lambda_{ij}^{(k)}\right)^{y_{ij}}\exp\left(-\lambda_{ij}^{(k)}\right)}{y_{ij}!}\Big]^{I(z_{ij}=k)}\exp\left\{-\frac{(\beta^{(k)}-m^{(k)})^2}{2(\sigma^2)^{(k)}}\right\} \tag{S6}$$

where $m^{(k)}$ is the mean and $\left(\sigma^{(k)}\right)^2$ is the variance for component $k$.

To update the latent variable, the probability of an observation belonging to each component is calculated

$$Pr(z_s|\gamma, z_t, \boldsymbol{y}, \vec{\boldsymbol{\beta}}^{(\boldsymbol{k})}) \propto e^{\gamma\sum_{s\sim t}\delta_{z_s z_t}} f(y_{i,j};\boldsymbol{\beta}^{(k)}) \tag{S7}$$

where $s$ is fragment pair $i$ and $j$, and $t$ is the neighbours set of $s$; ($i-1$, $i+1$, $j-1$, $j+1$) and $f^{(k)}(y_{ij};\vec{\boldsymbol{\beta}}^{(\boldsymbol{k})})$ is the likelihood of component $k$.

When the normalizing constant is introduced, equation S7 can be rewritten as

$$Pr(z|\gamma, z_t, \boldsymbol{y}, \vec{\boldsymbol{\beta}}^{(k)}) = \frac{e^{\gamma\sum_{s\sim t}\delta_{z_s z_t}}}{\sum_{z_s} e^{\gamma\sum_{s\sim t}\delta_{z_s z_t}}}\frac{f(y_{i,j};\boldsymbol{\beta}^{(k)})}{f(y_{i,j};\boldsymbol{\beta}^{(k)})} \tag{S8}$$

where $s$ is fragment pair $i$ and $j$, and $t$ is the neighbour(s).

The conditional probability of $\gamma$ in the Potts model is given as

$$Pr(\gamma|\boldsymbol{y},\boldsymbol{z},\vec{\boldsymbol{\beta}}) = \frac{\exp\{\gamma\sum_{s\sim t}\delta(z_s z_t)\}\pi_0(\gamma)}{\sum_{z_s}\exp\{\gamma\sum_{s\sim t}\delta(z_s z_t)\}\pi_0(\gamma)} \tag{S9}$$

---

**Algorithm S2** Metropolis-within-Gibbs sampler

---

**procedure**
  Initialization, select initial value,$\boldsymbol{z}^0, \boldsymbol{\gamma}^0, \boldsymbol{\beta}^0$;
  **repeat**
    **for** $i = 1$ to $n$, $j = 1$ to $n$ **do**
      Update $z_{ij}$ using (5.13)
    **end for**
    Update $\vec{\boldsymbol{\beta}}$ from posterior in (5.10)
    Update $\boldsymbol{\gamma}$ using Algorithm (0.1)
    Update $\boldsymbol{\tau}$ using (5.8)
  **until** enough MCMC steps have been simulated;

---

**Algorithm S1** ABC

**procedure**
  **repeat**
    Select the initial value of $\gamma_0$ from the prior.
    $m = 0$
    **for** $i = 1 : N$ **do**
      Compute a new $y^*$ from the Potts model
      Compute the distance $d(y, y^*)$
      Select epsilon using $1\%$ empirical quantile of $d(y, y^*)$
      **if** $d(y, y^*) < \epsilon$ **then**
        $\gamma_{m+1} = (\gamma_m)^*$
      **end if**
    **end for**
  **until** enough MCMC steps have been simulated;

The equations for $\boldsymbol{\beta}$'s and $\boldsymbol{z}$ as given above in equations S3, S4 and S7, is computationally easier to simulate using the Metropolis-Hastings-within-Gibbs sampler. Equation S9 is computationally intractable as the interaction parameter $\gamma$ involves the evaluation of the partition function and cannot be simulated directly using the Gibbs sampler or the Metropolis-Hastings sampler.

Algorithm $0.1$ shows the Approximate Bayesian Computation (ABC) approximate steps and Algorithm $0.2$ shows the Metropolis-within-Gibbs steps used in this paper to update our parameters.

### S2.2 Analysis

Table S1: Simulation results for normal priors. $\beta_0^{(1)}$ and $\beta_1^{(1)}$ are the Intercept and Distance parameters of the noise component, while $\beta_0^{(2)}$ and $\beta_1^{(2)}$ are the Intercept and Distance parameters of the signal components. In brackets we presented the $95\%$ credible intervals. For the fixed prior, we used $\beta_0^{(1)} \sim N(1, 1)$, $\beta_0^{(2)} \sim N(150, 10)$, $\beta_1^{(1)} \sim N(0.5, 0.5)$ and $\beta_1^{(2)} \sim N(1, 0.5)$.

| Parameters | True value | Posterior mean (fixed prior) | Posterior mean (empirical bayes method) |
|---|---|---|---|
| $\beta_0^{(1)}$ | 0.05 | 0.04 (0.01, 0.05) | 0.04 (0.01, 0.07) |
| $\beta_1^{(1)}$ | 0.2 | 0.21 (0.24, 0.29) | 0.22 (0.2, 0.27) |
| $\beta_0^{(2)}$ | 5.00 | 4.99 (4.99, 5.00) | 5.00 (4.98, 5.01 ) |
| $\beta_1^{(2)}$ | 2 | 1.98 (1.99, 2.04) | 2.02 (1.93, 2.08) |

Table S2: Simulation results for normal priors. $\beta_0^{(1)}$, $\beta_1^{(1)}$, $\beta_2^{(1)}$,$\beta_3^{(1)}$ and $\beta_4^{(1)}$ are the Intercept, Distance, GC-content, TEs and Accessibility parameters of the noise component, while $\beta_0^{(2)}$, $\beta_1^{(2)}$, $\beta_2^{(2)}$ and $\beta_3^{(2)}$ are the Intercept, Distance, GC-content, TEs and Accessibility parameters of the signal components. In bracket are the $90\%$ credible intervals.

| Parameters | True value | Posterior mean |
|---|---|---|
| $\beta_0^{(1)}$ (intercept) | 0.05 | 0.04 (0.05, 0.15) |
| $\beta_1^{(1)}$ (distance) | 0.2 | 0.18 (0.14, 0.24) |
| $\beta_2^{(1)}$ (GC-content) | 0.3 | 0.30 (0.25, 0.35) |
| $\beta_3^{(1)}$ (TEs) | 0.2 | 0.19 (0.14, 0.25) |
| $\beta_4^{(1)}$ (Accessibility) | 0.1 | 0.08 (0.03, 0.13) |
| $\beta_0^{(2)}$ (intercept) | 5 | 5.01 (4.96, 5.06) |
| $\beta_1^{(2)}$ (distance) | 2 | 2.01 (1.96, 2.06) |
| $\beta_2^{(2)}$ (GC-content) | 0.8 | 0.80 (0.75, 0.85) |
| $\beta_3^{(2)}$ (TEs) | 0.7 | 0.69 (0.65, 0.74) |
| $\beta_4^{(2)}$ (Accessibility) | 0.6 | 0.60 (0.55, 0.65) |

| Parameters | True value | Posterior mean (empirical Bayes method) |
|---|---|---|
| $\beta_0^{(1)}$ | 0.05 | 0.06 (0.02 0.11) |
| $\beta_1^{(1)}$ | 0.2 | 0.20 (0.15 0.24) |
| $\beta_0^{(2)}$ | 5 | 5.00 (4.97 5.03) |
| $\beta_1^{(2)}$ | 2 | 2.02 (1.83 2.21) |

Table S3: Simulation results for normal priors when the proportion of signal = noise. $\beta_0^{(1)}$ and $\beta_1^{(1)}$ are the Intercept and Distance parameters of the noise component, while $\beta_0^{(2)}$ and $\beta_1^{(2)}$ are the Intercept and Distance parameters of the signal components. In bracket are the $90\%$ credible intervals.

| Parameters | True value | Posterior mean (empirical Bayes method) |
|---|---|---|
| $\beta_0^{(1)}$ | 0.05 | 0.09 (0.03 0.19) |
| $\beta_1^{(1)}$ | 0.2 | 0.16 (0.11 0.20) |
| $\beta_0^{(2)}$ | 5 | 5.00 (4.98) 5.02) |
| $\beta_1^{(2)}$ | 2 | 1.96 (1.74 2.21) |

Table S4: Simulation results for normal priors when the proportion of noise = 0.3 and the proportion of signal = 0.7. $\beta_0^{(1)}$ and $\beta_1^{(1)}$ are the Intercept and Distance parameters of the noise component, while $\beta_0^{(2)}$ and $\beta_1^{(2)}$ are the Intercept and Distance parameters of the signal components. In bracket are the $90\%$ credible intervals.

Table S5: Posterior means of our estimated $\beta$s for both noise and signal components. The 95% credible intervals are shown inside the brackets.

| Parameters | Posterior mean noise (95% intervals) | Posterior mean signal (95% intervals) |
|---|---|---|
| $\beta_0$ (intercept) | -84.26 (-85.26, -83.29) | 12.24 (11.26, 13.22) |
| $\beta_1$ (distance) | -10.10 (-11.08, -9.12) | -0.92 (-1.90, -0.06) |
| $\beta_2$ (GC-content) | 0.35 (-0.64, 1.34) | 4.38 (3.40, 5.37) |
| $\beta_3$ (TEs) | -0.73 (-1.73, 0.23) | -0.02 (-1.00, 0.96) |
| $\beta_4$ (Accessibility) | -3.51 (-4.49, -2.52) | 0.07 (-0.91, 1.05) |

As mentioned in the main text, we simulated only one source of bias (distance) to see how our method performed to different proportions of noise and signal due to the computation time. Table S3 shows the result of the simulation study when the proportion of noise and signal are the same. In Table S3, when the proportion of noise and signal in the simulated data are the same, we can see that our method using the empirical Bayes method adequately estimated the true value of the parameters in our simulated data. Table S4 shows the result of the simulation study when the proportion of noise is less than that of signal proportion. We can see that our method using the empirical Bayes method as shown in Table S4 adequately estimated the true value of the parameters in our simulated data. The $90\%$ credible intervals in both Tables S3 and S4 are all significant.

From Table S5, we can see that the credible intervals for $\beta_2$ and $\beta_3$ (GC-content and TEs) are not significant at $95\%$ credible intervals for the noise component. For the signal component, $\beta_3$ and $\beta_4$ (TEs and Accessibility) are not significant. Due to about half of our covariates not being significant, when we set our credible interval as $95\%$, we instead use $90\%$ for our analysis.

### S2.3 Hi-C Data analysis with a two components model

Table S6: Posterior means of our estimated $\beta$s shown in equation 2 for both noise and signal components. The 90% credible intervals are shown inside the brackets.

| Parameters | Posterior mean noise (90% intervals) | Posterior mean signal (90% intervals) |
|---|---|---|
| $\beta_0$ (intercept) | -84.26 (-84.86, -83.73) | 12.24 (11.86, 12.60) |
| $\beta_1$ (distance) | -10.10 (-10.14, -10.06) | -0.92 (-0.94, -0.90) |
| $\beta_2$ (GC-content) | 0.35 (0.34, 0.35) | 4.38 (3.71, 5.07) |
| $\beta_3$ (TEs) | -0.73 (-0.79, -0.70) | -0.02 (-0.07, 0.03) |
| $\beta_4$ (Accessibility) | -3.51 (-3.55, –3.46) | 0.07 (0.02, 0.10) |

To better understand the contributions of the different components, we investigated the posterior means of our estimated $\beta$s for both the noise and signal components (see Table S6).

For the signal component, we noticed that the posterior means of the coefficient of the distance and TEs ($\beta_1$ and $\beta_3$) are negative values. While the posterior means of the intercept, GC-content and DNA accessibility ($\beta_0$, $\beta_2$ and $\beta_4$) are all positive values. In addition, the credible intervals

Table S7: Posterior means of our estimated $\beta$s as shown in equation 2 for noise, signal and false signal components. The 90% credible intervals are shown inside the brackets.

| Parameters | Posterior mean (noise) | Posterior mean (signal) | Posterior mean (false signal) |
|---|---|---|---|
| $\beta_0$ (intercept) | -84.00 (-84.80, -83.90) | 13.06 (12.86, 13.61) | 499.34 (498.67, 500.00) |
| $\beta_1$ (distance) | -10.05 (-10.13, -10.04) | -0.90 (-0.91, -0.89) | -64.16 (-64.14, -64.01) |
| $\beta_2$ (GC content) | 0.34 (0.34, 0.35) | 0.36 (0.35, 0.37) | 0.30 (0.15, 0.52) |
| $\beta_3$ (TEs) | -0.76 (-0.80, -0.72) | -0.10 (-0.15, -0.05) | 0.54 (-0.33, 0.80) |
| $\beta_4$ (Accessibility) | -3.54 (-3.56, -3.48) | 0.15 (0.12, 0.19) | -0.70 (-0.89, -0.27) |

for distance ($\beta_1$) is significant and the negative posterior mean indicates that as the distance of two fragments increases, the average of their signal interaction decreases as well. Similarly for the other significant parameters, $\beta_2$ (GC content) and $\beta_4$ (DNA accessibility), their positive values for posterior means and credible intervals indicates that as GC-content and level of DNA accessibility increases the average of the signal interaction increases as well. In other words, our results indicate that there is a small impact of DNA accessibility on the Hi-C results, where regions with higher DNA accessibility are retrieved more often than regions in dense chromatin, but this bias is small. However for $\beta_3$ of the signal component, the credible intervals is not significant as it is having $0$ in-between. Altogether, our results show that genomic distance between pairs of loci and the GC-content are the most significant sources of bias in our Hi-C data (Table S6).

### S2.4 Genome wide analysis of *Drosophila* Kc167 cells

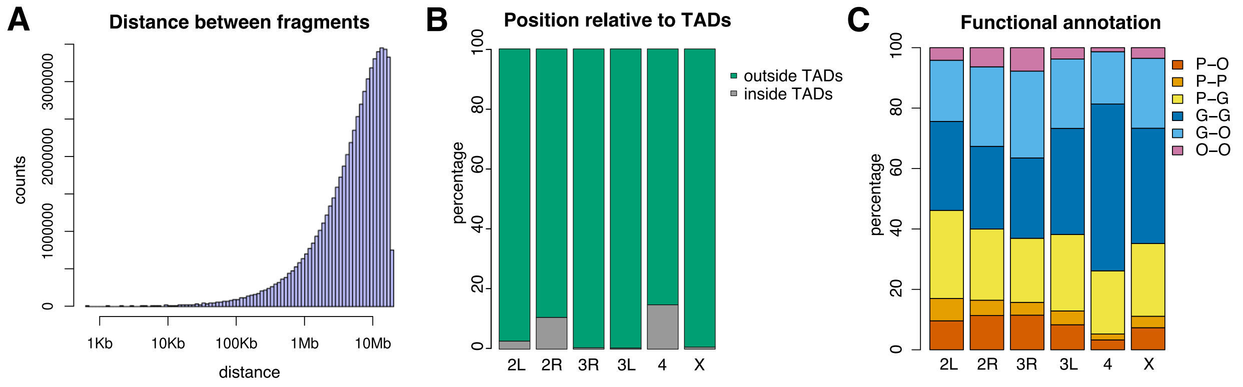


Figure S2: *Genome wide significant interactions in Drosophila Kc167 cells.* (A) Distribution of the distance between the two fragments for all significant interactions. (B) classification of significant interactions as either outside TADs when the two fragments are located in different TADs or inside TADs when the two fragments are located in the same TAD. (C) Percentage of significant interactions that have promoters at one of the fragments. We consider the cases of: (P-P) both fragments contain promoters, (P-O) only one fragment contains promoters and (O-O) none of the fragments contain any promoter.

### S2.5 Data analysis of Hi-C and micro-C in human ES cells

Table S8: Posterior means of our estimated $\beta$s as shown in equation 2 for noise, signal and false signal components of human Chromosome 8, region 60M : 70M for data generated using the Hi-C method. The 95% credible intervals are shown inside the brackets.

| Parameters | Posterior mean (noise) | Posterior mean (signal) | Posterior mean (false signal) |
|---|---|---|---|
| $\beta_0$ (intercept) | 1.06 (0.79, 1.49) | 5.84 (5.67, 5.99) | 5.69 (5.53, 5.79) |
| $\beta_1$ (distance) | 0.15 (0.14, 0.17) | -0.99 (-1.00, -0.98) | -0.99 (-1.01, -0.98) |
| $\beta_2$ (GC content) | 0.32 (0.32, 0.33) | -0.58 (-0.84, -0.21) | -0.07 (-0.29, 0.18) |
| $\beta_3$ (TEs) | 10.00 (9.97, 10.03) | -0.09 (-0.10, -0.07) | -0.1 (-0.13, -0.09) |
| $\beta_4$ (Accessibility) | 0.38 (0.35, 0.40) | 0.95 (0.92, 0.98) | 1.10 (0.97, 1.19) |

Table S9: Posterior means of our estimated $\beta$s as shown in equation 2 for noise, signal and false signal components of human Chromosome 8, region 60M : 70M for data generated using the micro-C method. The 95% credible intervals are shown inside the brackets.

| Parameters | Posterior mean (noise) | Posterior mean (signal) | Posterior mean (false signal) |
|---|---|---|---|
| $\beta_0$ (intercept) | 0.98 (0.55, 1.58) | 7.91 (7.73, 8.09) | 8.33 (8.08, 8.53) |
| $\beta_1$ (distance) | 0.15 (0.12, 0.17) | -1.40 (-1.43, -1.38) | -1.41 (-1.43, -1.38) |
| $\beta_2$ (GC content) | 0.32 (0.32, 0.33) | 1.60 (1.14, 1.96) | 0.30 (-0.36, 0.98) |
| $\beta_3$ (TEs) | 10.00 (9.97, 10.04) | -0.37 (-0.41, -0.33) | -0.37 (-0.40, -0.34) |
| $\beta_4$ (Accessibility) | 0.38 (0.35, 0.41) | 1.77 (1.58, 1.94) | 1.81 (1.71, 1.86) |

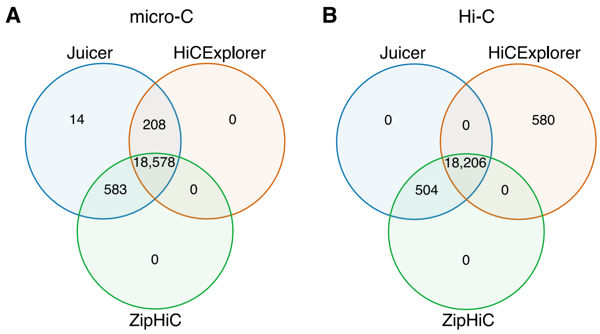


Figure S3: *Comparison between ZipHiC, HiCExplorer and Juicer on human data.*. (A) We considered the region 60-70Mb of the human chromosome 8 and data from (A) micro-C and (B) Hi-C in human ES cells.
